# Early immune response in mice immunized with a semi-split inactivated vaccine against SARS-CoV-2 containing S protein-free particles and subunit S protein

**DOI:** 10.1101/2020.11.03.366641

**Authors:** Marek Petráš, Petr Lesný, Jan Musil, Radomíra Limberková, Alžběta Pátíková, Milan Jirsa, Daniel Krsek, Pavel Březovský, Abhishek Koladiya, Šárka Vaníková, Barbora Macková, Dagmar Jírová, Matyáš Krijt, Ivana Králová Lesná, Věra Adámková

## Abstract

The development of a vaccine against COVID-19 is a hot topic for many research laboratories all over the world. Our aim was to design a semi-split inactivated vaccine offering a wide range of multi-epitope determinants important for the immune system including not only the spike (S) protein but also the envelope, membrane and nucleocapsid proteins. We designed a semi-split vaccine prototype consisting of S protein-depleted viral particles and free S protein. Next, we investigated its immunogenic potential in BALB/c mice. The animals were immunized intradermally or intramuscularly with the dose adjusted with buffer or addition of aluminum hydroxide, respectively. The antibody response was evaluated by plasma analysis at 7 days after the first or second dose. The immune cell response was studied by flow cytometry analysis of splenocytes. The data showed a very early onset of both S protein-specific antibodies and virus-neutralizing antibodies at 90% inhibition regardless of the route of vaccine administration. However, significantly higher levels of neutralizing antibodies were detected in the intradermally (geometric mean titer - GMT of 7.8 ± 1.4) than in the intramuscularly immunized mice (GMT of 6.2 ± 1.5). In accordance with this, stimulation of cellular immunity by the semi-split vaccine was suggested by elevated levels of B and T lymphocyte subpopulations in the murine spleens. These responses were more predominant in the intradermally immunized mice compared with the intramuscular route of administration. The upward trend in the levels of plasmablasts, memory B cells, Th1 and Th2 lymphocytes, including follicular helper T cells, was confirmed even in mice receiving the vaccine intradermally at a dose of 0.5 μg.

We demonstrated that the semi-split vaccine is capable of eliciting both humoral and cellular immunity early after vaccination. Our prototype thus represents a promising step toward the development of an efficient anti-COVID-19 vaccine for human use.

## Introduction

The COVID19 pandemic has made the development of a vaccine an emergency priority. Hectic research was enabled by recent major advances in sequencing, protein structure identification, and epitope mapping that have suggested the spike (S) protein of SARS-CoV-2 virus as a top candidate for vaccine construction to elicit effective antibodies [Hoffmann 2020, Robbiani 2020] which was why vaccine researchers all over the world focused exclusively on this particular immunogen. The first vaccines in clinical trials are designed not only as traditional ones (i.e., inactivated or live) but also as RNA vaccines not used in human clinical research to date. In fact, except for wholly inactivated vaccines that include other important antigens, all vaccines currently being tested contain only the S protein [Xia 2020, Zhang 2020, Pu 2020].

S protein-based vaccines are designed as subunit ones [Keech 2020]. Alternatively, the protein is expressed in host cells from non-replicating (defective) viral vectors currently using simian or human adenovirus type 5 or 26 [Zhu 2020, Logunov 2020, Folegatti 2020, Sadoff 2020] or from mRNA encapsulated in lipid nanoparticles [Jackson 2020, Sahin 2020, Mulligan 2020, Walsh 2020]. These vaccines clearly exhibited a great immunogenic potential diminished by an increased frequency of adverse events in the early trials. Conversely, inactivated virion vaccines have consistently showed reasonable immunogenicity with a very good safety profile.

Recent computational approaches have suggested that the envelope (E), membrane (M) and nucleocapsid (N) proteins should be included in multi-epitope vaccines as these proteins exhibit conserved sequence features with a high affinity for the major histocompatibility complex (MHC) [Ghafouri 2020, Khairkhah 2020, Noorimotlagh 2020, Sarkar 2020].

Unlike the worldwide commercial efforts to develop efficient and safe vaccines, we sought to construct an emergency vaccine under a project referred to as the “Emergent Coronavirus Vaccine” (EmCoVax). The principal aim of the first phase was to develop and test the process of vaccine production. We decided to design a semi-split inactivated vaccine containing separately S protein and S protein-free viral particles. Early immunogenicity was subsequently investigated in a mouse model within two weeks of immunization to test the very rationale of this type of vaccine.

## Materials and Methods

### Viruses and tissue cultures cells

SARS-CoV-2 (strain 951) was selected from the virus archive of the National Institute of Public Health of the Czech Republic. Virus stock was produced in Vero E6 cells obtained from the American Type Culture Collection (VERO 76, ATCC® CRL-1587™) under serum-free conditions using OptiPro medium (Life Technologies Europe BV, Bleiswijk, the Netherlands).

### Electron microscopic examination

Approximately 500 μL of inactivated SARS-CoV-2 suspension was placed on a parafilm with attached two formvar/carbon-coated TEM grids pretreated by 1% Alcian blue to increase hydrophilicity and allowed to stand for 20 minutes. The first grid was stained with 2% aqueous solution of ammonium molybdate and the second one with 2% aqueous solution of uranyl acetate. After removing the residual fluid using filter paper, the grids were observed under a transmission electron microscope (Hitachi HT-7800, Hitachi, Tokyo, Japan).

### Virus titer determination

The infectious virus titer of SARS-CoV was determined by the tissue culture infectious dose 50% (TCID_50_) assay. In brief, serial 10-fold dilutions of virus-containing samples were inoculated onto confluent Vero E6 monolayers in 96-well plates. After incubation for 5 days at 37°C in a CO_2_ incubator, the cells were assessed for the presence of a cytopathic effect (CPE) by means of an inverted optical microscope. TCID_50_ was determined by a 50% reduction in CPE using the Reed-Muench formula [Ramakrishnan 2016].

### Determination of S protein content

The concentration of the S protein was determined with an ELISA kit (SARS-CoV-2 Spike ELISA kit, Sino Biological Inc., Beijing, China). A monoclonal antibody specific for the S protein of SARS-CoV-2 was pre-coated onto well plates. Standards and samples were added to the wells and the S protein present in the sample was bound by the immobilized antibody. After incubation, the wells were washed and a horseradish peroxidase-conjugated anti-S antibody added, producing a sandwich complex. After washing to remove the unbound antibody, a tetramethylbenzidine (TMB) substrate solution was loaded. The reaction was stopped by the addition of a stop solution and the color intensity was measured at 450 nm. To determine the S protein concentration in the sample, a standard curve was generated at concentrations of the working standard, within a range of 2,500-10,000 pg/mL of the S protein.

### Virus neutralization assay (VNT_90_)

Mouse plasma were tested for functional antibodies against SARS-CoV-2 with the standard virus neutralization test. Prior to the test, each plasma sample was heated at 56°C for 30 minutes. Serial 2-fold dilutions of the plasma from 1:2 to 1:32 were subsequently incubated with 100 TCID_50_ of the vaccine of the SARS-CoV-2 virus strain for 2 hours and added to VERO E6 cell monolayers. The characteristic CPE was read on Day 5. Neutralizing antibody titers were expressed as the reciprocal of the last plasma dilution that completely inhibited CPE in 90% of the wells using the Reed-Muench formula [Tseng 2012, Zakhartchouk 2005, Spruth 2006].

### ELISA antibodies assay

Specific anti-SARS-CoV-2 IgG antibodies in mice were detected with a commercial ELISA kit (Human Anti-2019 nCoV(S) IgG ELISA Kit, FineTest, Wuhan, China) where the purified horseradish peroxidase (HRP)-conjugated anti-mouse IgG detection antibodies were replaced by HRP-conjugated AffiniPure goat anti-mouse IgG (Wuhan Fine Biotech Co., Ltd. China) diluted with an antibody dilution buffer at 1:1,000 according to the anti-mouse IgG manufacturer’s instructions. ELISA plates were pre-coated with a recombinant 2019-nCoV spike protein. The assay provided semiquantitative results by calculating the ratio of the optical densities (ODs) of the plasma samples over the negative control OD. Since negative mouse plasma was unavailable, dilutions of plasma samples and their ODs were log-transformed, and the intercept of the regression straight line on the dilution axis represented the log-titer of antibodies. To avoid overestimation of the titer, its value was assessed as an antilogarithm of the lower limit of the 95% confidence interval of this intercept.

### Flow cytometry analysis

Samples preparation was carried out according to standard protocols (see Supplement). Briefly, spleens from mice were harvested, pooled in each group and dissociated using the gentleMACS Octo dissociator (Miltenyi Biotec, Bergish Gladbach, Germany) according to the manufacturer’s protocol for preparation of single-cell suspensions from the murine spleen without enzymatic treatment. The resultant single-cell suspension was further purified using the standard Ficoll density gradient [Cossarizza 2019]. Purified cells were subsequently resuspended in CryroStor medium (MilliporeSigma, St. Louis, MO) and stored in liquid nitrogen until analysis. A 24-color spectral cytometry panel was designed to characterize T and B cell differentiation and activation status. Cells were stained using standard protocols and acquired using the Aurora spectral cytometer (Cytek Biosciences, Fremont, CA). Data analysis was carried out using manual gating or unsupervised clustering analysis (see Supplement).

### Animal experiment

All animals were maintained in a conventional animal facility in a 12/12 light/dark regimen, controlled temperature and free access to pelleted food and water. All protocols were approved by the Animal Care Committee of the Institute for Clinical and Experimental Medicine and the Ministry of Health of the Czech Republic.

A total of 52 BALB/c mice (Velaz, Czech Republic) were immunized with vaccine (40 mice) or placebo (12 mice) at age 9-10 weeks. The vaccinated mice were divided into 8 groups of 5 mice immunized with one or two intramuscular or intradermal doses containing 0.5 or 1.5 μg of the S protein. A total of 12 mice, i.e., 4 groups of 3 mice with one or two intradermal or intramuscular doses, received placebo (0.9% NaCl solution). The volume of each dose, whether of vaccine or placebo, was consistently 10 μL. All mice received the first dose on Day 0, with half of them immunized with the second dose on Day 7.

Disposable insulin syringes with integrated needle (BD Micro-fine plus; 0.3 ml, Becton, Dickinson, NJ) were used for both routes of administration. While the intramuscular dose was applied into the right caudal thigh muscle, the intradermal one was applied into the auricle under the guidance of a microscope.

One week after the first or second dose, the mice were anesthetized with isoflurane and blood was intracardially withdrawn using a 2 ml syringe, transferred to anticoagulant tubes (K_2_EDTA), and mixed. Whole blood was centrifuged (3300 rpm, 10 minutes, 4°C), plasma aliquoted into 100 μl microtubes and frozen at −80° C. The spleens were collected in chilled RPMI medium on ice and transported to laboratory within 4 hours for flow cytometry examination.

### Statistical analysis

Anti-S specific IgG antibodies as well as virus-neutralizing antibodies were expressed as the titer. The value had to be log-transformed to pass a normality test (D’Agostino & Pearson or Shapiro-Wilk test) followed by a parametric t-test or analysis of variance (ANOVA).

As the lymphocyte populations were measured from pooled spleens of each group, the traditional statistical approach could not be employed; hence, the change in lymphocyte subsets using linear regression was determined. If the slope of lines exhibited values significantly different from null, then the observed change was considered to be confirmed. Merged lymphocyte subpopulations were analyzed with parametric tests after log-transformation to pass a normality test (Shapiro-Wilk test). The power of each test was insufficient since the sample size was small. A >80% power of the test was only achieved when comparing the geometric mean titers of antibodies between immunized and unimmunized mice.

The correlation between IgG antibodies and virus-neutralizing antibodies was assessed with the Pearson correlation coefficient after the log-transformation of their values.

Continuous data were summarized using standard descriptive statistics, i.e., median including range or geometric mean with standard deviation or 95% confidence interval. All tests were two-tailed, and the level of significance was set at 0.05. Statistical analyses were performed using Prism 8 (GraphPad Software, Inc., San Diego, CA) and STATA version 16 software (StatCorp, College Station, TX).

## Results

### Course of the culture, inactivation and purification process

To obtain a viral stock for vaccine production, the virus strain 951 was first purified by 4 passages in VERO E6 cells. Subsequently, another 8 passages were performed to generate a sufficient stock of virus for the production of the master seed as the source of a virus bank for vaccine production through the seed lot system. Growth kinetics analysis of the second passage showed a sufficient virus replication rate reaching a titer of 6.0-6.5 log_10_ TCID_50_/mL in the supernatant within 2-8 days (Figure 1). The peak titers of 6.5 and 7.5 log_10_ TCID_50_/mL from the supernatant and lysate, respectively, were at 56 hours post-infection. Moreover, multiplicities of infection (MOI) of 0.001-0.0001 at a culture temperature of 37°C and 5% CO_2_ were confirmed.

**Figure 1:**
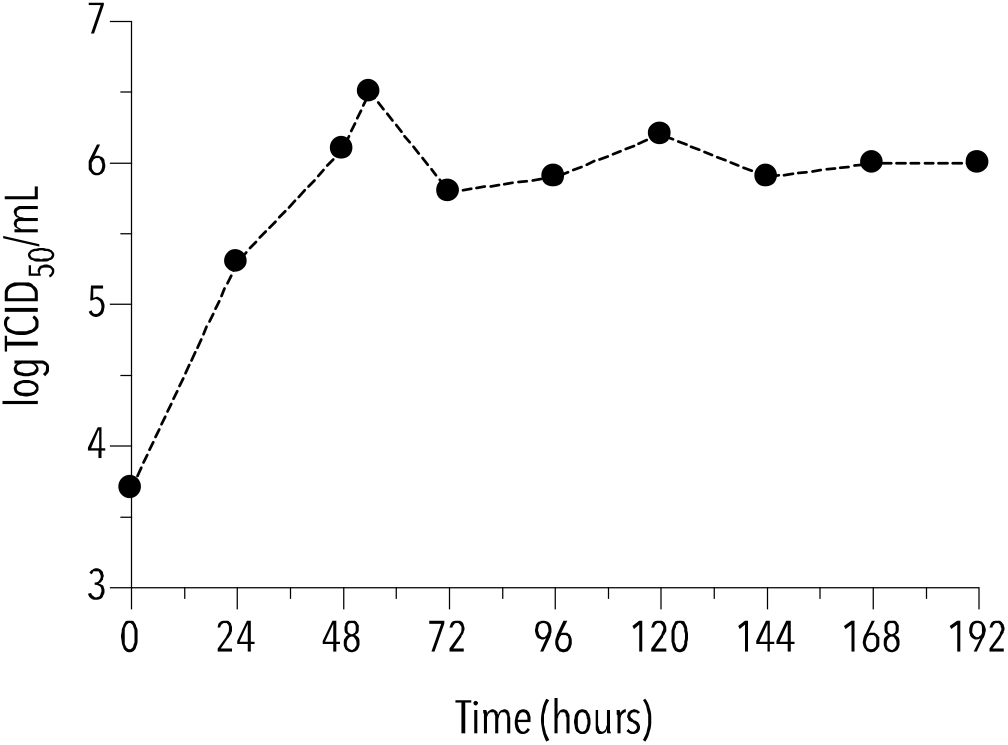
Growth kinetics analysis of viral titer obtained from the supernatant of the culture (TCID_50_ … tissue culture infectious dose 50%)

Confluent monolayers of Vero E6 cells, grown in OptiPro serum-free medium (Life Technologies Europe BV, the Netherlands) with 10% fetal bovine serum (FBS), were infected by the strain from the second passage of the stock and incubated in a serum-free medium at 37°C and 5% CO_2_ for 56 hours. The first viral suspension of 225 mL was obtained by centrifugation of the supernatant harvest at 300 g for 5 minutes. The second suspension was harvested from infected cells by adding an approx. 90 mL of medium, overnight freezing and thawing at −20°C followed by centrifugation under the same conditions. Both live viral suspensions containing viral particles with typical corona spikes as documented by electron-microscopic inspection (Figure 2A) were stored at 4°C for at least 11 days.

**Figure 2:**
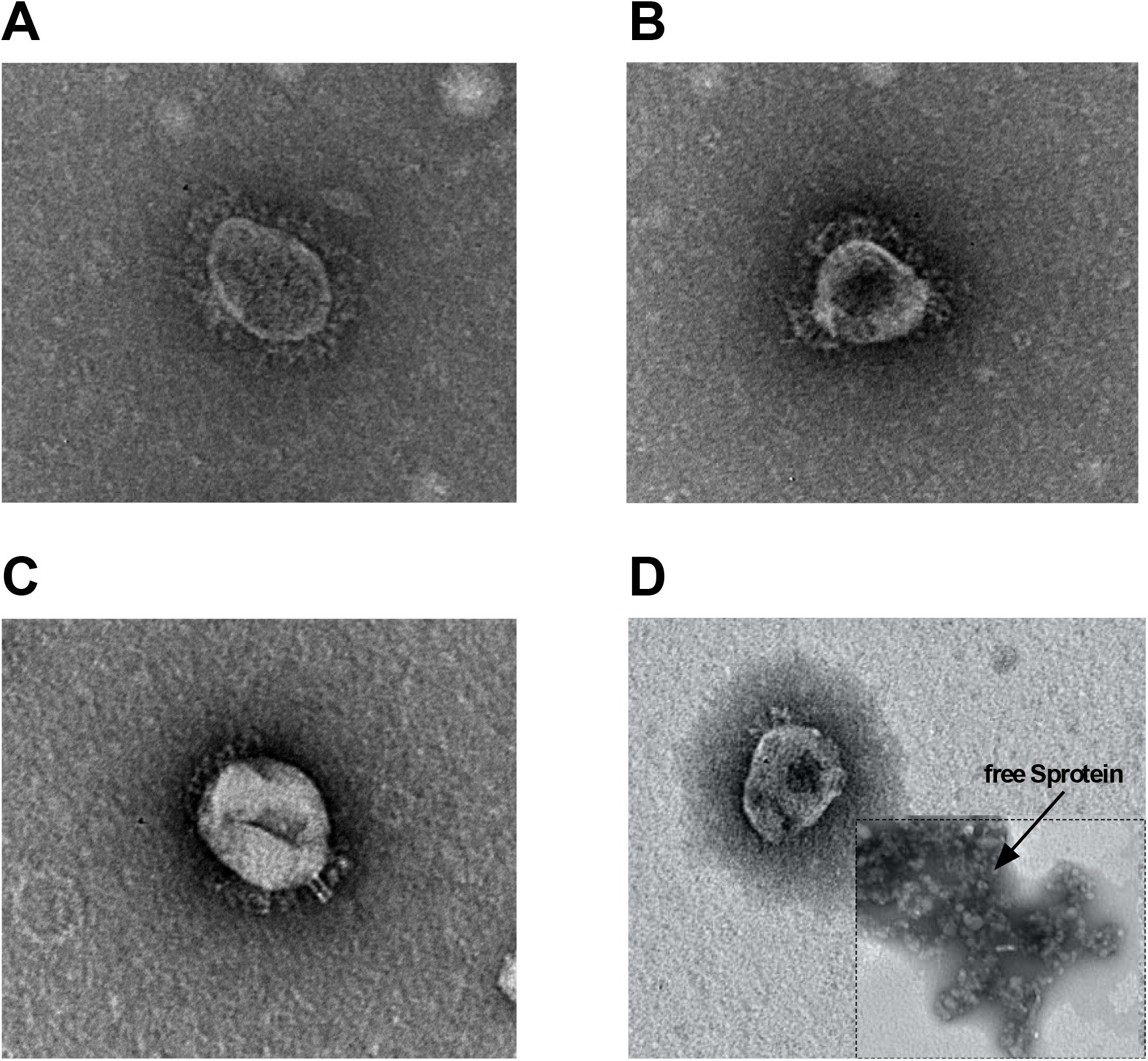
Electron micrograph: A – virus particle after culture; B – virus particle after the 1^st^ inactivation; C – virus particle after the 2^nd^ inactivation; D – virus particle and S protein after purification

The suspensions were subsequently inactivated by adding beta-propiolactone diluted 1:2,000, and continuously shaken up at a temperature of 4°C for 48 hours. The beta-propiolactone hydrolysis at 37°C lasted 2 hours. Inactivated suspensions were centrifuged and the supernatants were stored at 4°C for one week. The second inactivation was performed with beta-propiolactone diluted 1:1,000 at 4°C for 66 hours. After that, the pellets were centrifuged at 1,800 g for 10 minutes and both inactivated suspensions were subsequently placed in water baths to hydrolyze beta-propiolactone with continuous stirring at 37°C for 2 hours with the temperature gradually decreasing to 27°C overnight.

The inactivated viral suspension contained of 0.5 μg/mL of the S protein for the supernatant harvest and 4.9 μg/mL for the lysate harvest. While the first inactivation did not influence the virion structure, the second one showed partial changes, i.e., complete and incomplete particles with disrupted S protein (Figure 2B and 2C).

The inactivation including hydrolysis decreased the pH to 6.0-6.5 that disrupted S protein binding to the viral envelope as documented by a record from the electron microscope (Figure 2C). Both suspensions were stored at 4°C for 3 days to be followed by vaccine purification and thickening to obtain 3 μg/10 μL the for harvested supernatant and 1 μg/10 μL for the harvested lysate. This was achieved by a multiple ultra/diafiltration process utilizing Amicon#x00AE; Ultra centrifugal filter units (Amicon Ultracel) with a volume of 15 mL and a membrane molecular weight cutoff (MWCO) of 100 kDa (Amicon® Ultra-15 Centrifugal Filter Unit – 100 kDa cutoff, Merck Millipore Ltd., Darmstadt, Germany). The suspensions were centrifuged several times in Amicon Ultracel at 1,000 g for at least 5 minutes to exchange the culture medium and to remove cellular debris and low-molecular substances. The suspension was washed several times with phosphate buffer saline (PBS) of pH 7.4.

Inactivation of virus infectivity was confirmed by assaying absent CPE in Vero E6 cells within a dilution range of 10^−1^ to 10^−4^.

The result of purification and thickening were suspensions containing both virion particles free of the spike and separated S protein as documented by electron microscopy (Figure 2D). The vaccine dose for the test of immunogenicity in mice was adjusted in PBS to concentrations of 0.5 μg/10 μL for the harvested lysate and 1.5 μg/10 μL for the harvested supernatant. Since two different routes of administration, i.e., intradermal (ID) or intramuscular (IM) one, were anticipated, the IM doses were prepared by dilution in a 0.05% solution of aluminum hydroxide wet gel (Alhydrogel adjuvant 2%, InvivoGen) so the final concentrations were identical for either route of administration.

### Safety of immunization

No pathological manifestations were observed in any mouse during post-immunization follow-up as were not any reductions in feed or water intake. Behavioral manifestations and weight gain were physiological. No efflorescence or other pathological reactions were found at the site of inoculation.

All body surfaces and orifices of all animals were free of any pathology. Mild splenomegaly was observed in the spleen of two animals immunized intradermally suggesting a cellular immune response to vaccination. Other internal organs were free of signs of inflammation or other pathologies. Likewise, no pathological activation of the immune system described during infection with the wild virus was seen in our project.

### Immunogenicity in mice

A single dose administered intradermally or intramuscularly showed significantly increased anti-S IgG antibodies compared with mice receiving placebo. Although the rise in anti-S IgG antibody levels was small, it was proportional to a short post-vaccination time (Figure 3). Unfortunately, it was not possible to evaluate the statistical significance among different groups categorized by dose size and route of administration since the sample size was too small to achieve enough statistical power. Regarding both routes of administration, one can expect a somewhat earlier response in mice immunized with a lower antigen content no matter whether one or two doses were administered.

**Figure 3:**
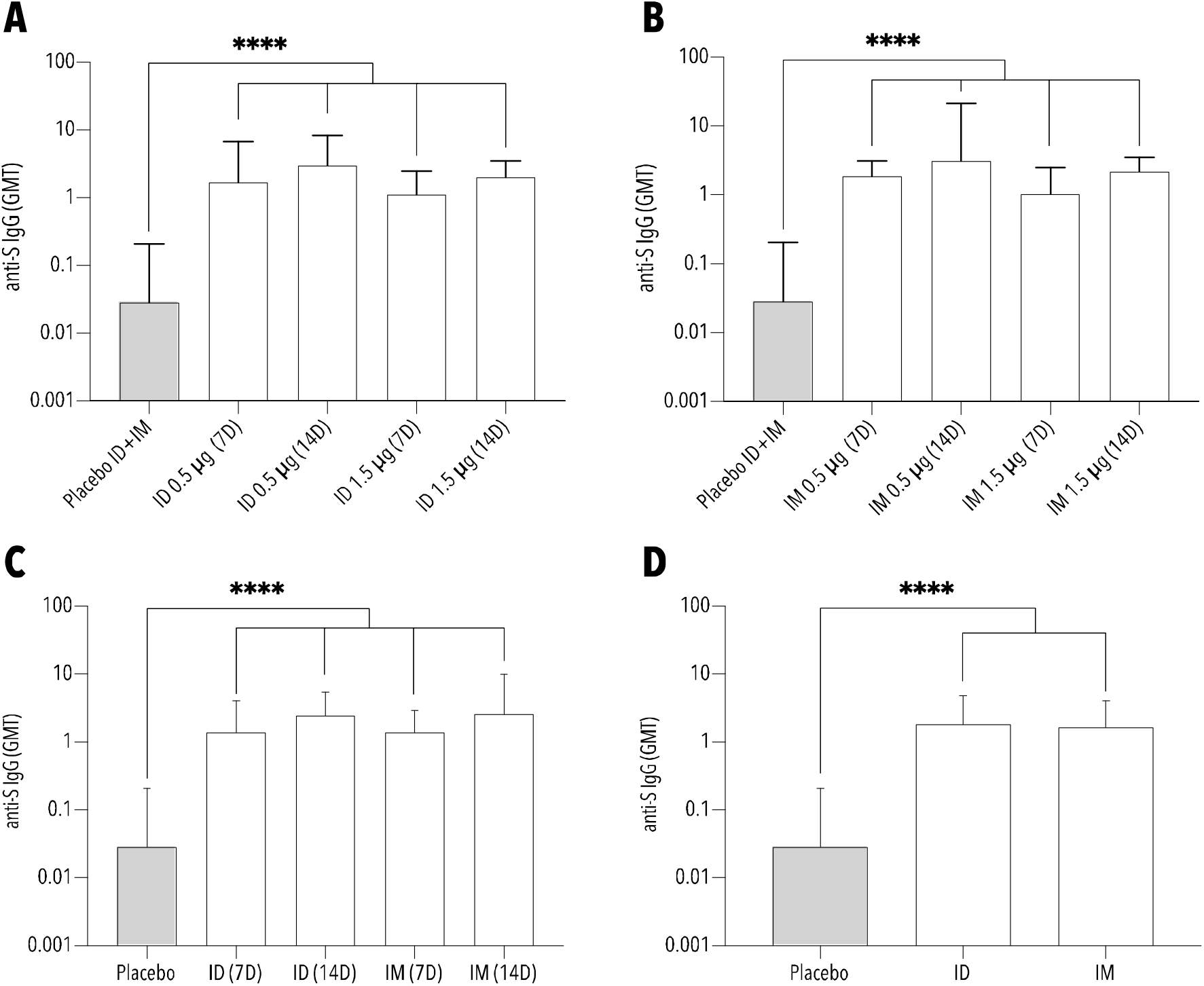
Geometric mean and geometric standard deviation of titer of anti-S IgG antibodies in immunized and unimmunized mice; A – groups of mice immunized intradermally with one (7 days) or two doses (14 days) of 0.5 and 1.5 μg; B – groups of mice immunized intramuscularly with one (7 days) or two doses (14 days) of 0.5 and 1.5 μg; C – groups of mice immunized intradermally and intramuscularly with one (7 days) or two doses (14 days); D – groups of mice immunized intradermally or intramuscularly irrespective of the size and number of doses (ID … intradermal administration; IM … intramuscular administration; 7D … 7 days; 14D … 14 days; GMT … geometric mean titer; anti-S … S protein antibodies; **** … p<0.0001)

Although the levels of anti-S IgG antibodies increased at 7 days after two-dose immunization, no statistical significance was found compared with the levels observed 7 days after the first dose (Figure 3). The same outcome was confirmed in both groups assessed according to the route of administration regardless of the antigen amount.

Functional antibodies determined as virus-neutralizing antibodies with 90% protective activity showed a significant increase at 7 days after the first or second dose independently of the route of administration (Figure 4). Although the second dose repeatedly increased the geometric mean of titers (GMT) of VNT_90_ antibodies independently of dose size or route of administration, the rate of growth was higher in mice immunized intradermally or intramuscularly with a dose of 1.5 μg. Despite this, the rise was not quite significant (p=0.08). Irrespective of the size dose, the GMT of neutralizing antibodies induced by the first intradermal dose was 6.8±1.3 and was significantly higher than that in the mice immunized intramuscularly, i.e., 5.1±1.4. While the second ID dose contributed to a non-significant increase in virus-neutralizing antibodies (GMT = 8.8±1.3), the second IM dose significantly increased geometric mean of neutralizing antibodies to 7.7±1.4 (p<0.05).

**Figure 4:**
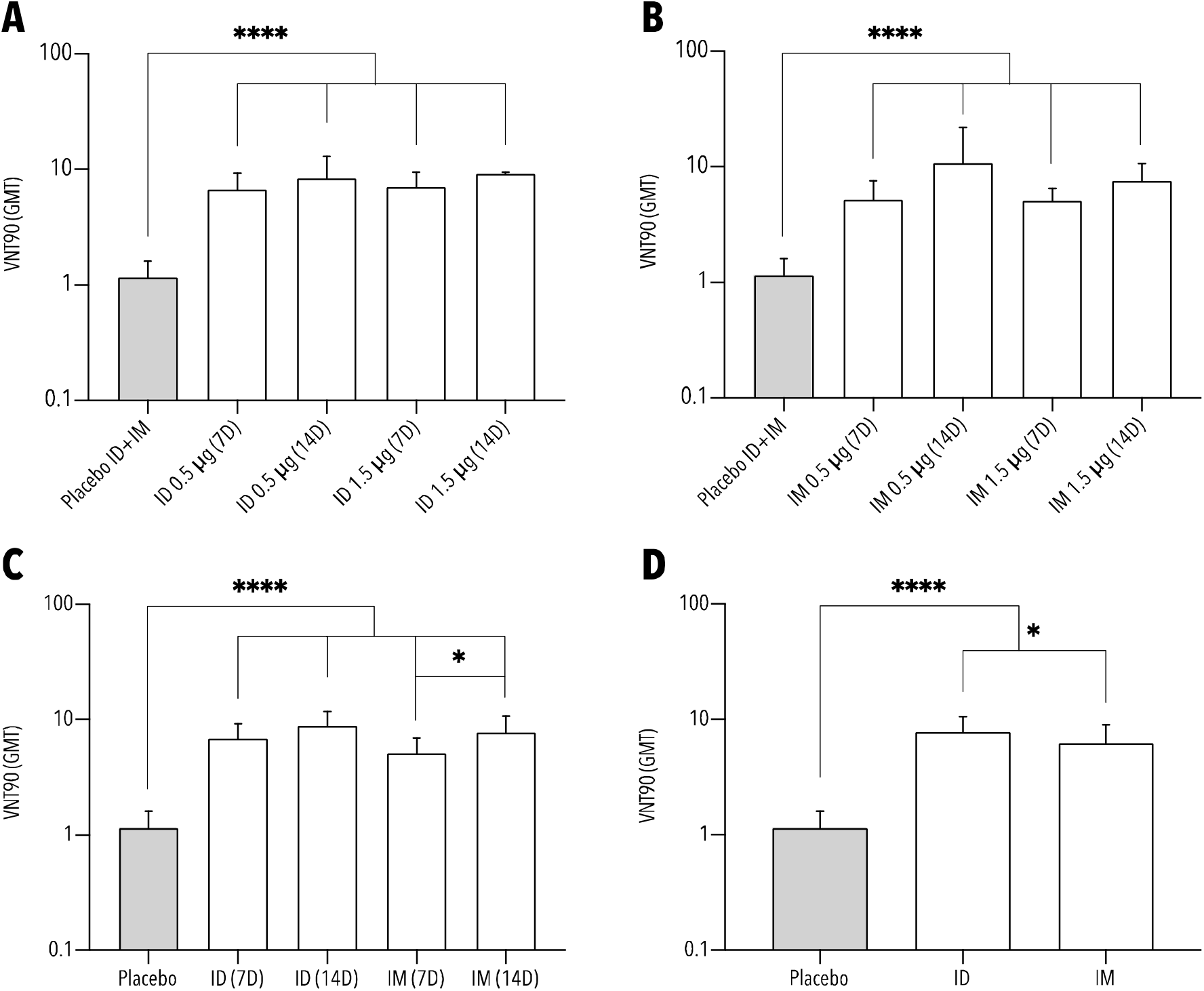
Geometric mean and geometric standard deviation of titer of VNT_90_ antibodies in immunized and unimmunized mice; A – groups of mice immunized intradermally with one (7 days) or two doses (14 days) of 0.5 and 1.5 μg; B – groups of mice immunized intramuscularly with one (7 days) or two doses (14 days) of 0.5 and 1.5 μg; C – groups of mice immunized intradermally and intramuscularly with one (7 days) or two doses (14 days); D – groups of mice immunized intradermally or intramuscularly irrespective of the size and number of doses (ID … intradermal administration; IM … intramuscular administration; 7D … 7 days; 14D … 14 days; GMT … geometric mean titer; VNT_90_ … virus-neutralizing titer inhibiting 90% of infectious dose; * … p<0.05; **** … p<0.0001)

Moreover, a good correlation between anti-S IgG and VNT_90_ titers of antibodies was documented, with a Pearson correlation coefficient of 0.86 (Figure 5).

**Figure 5:**
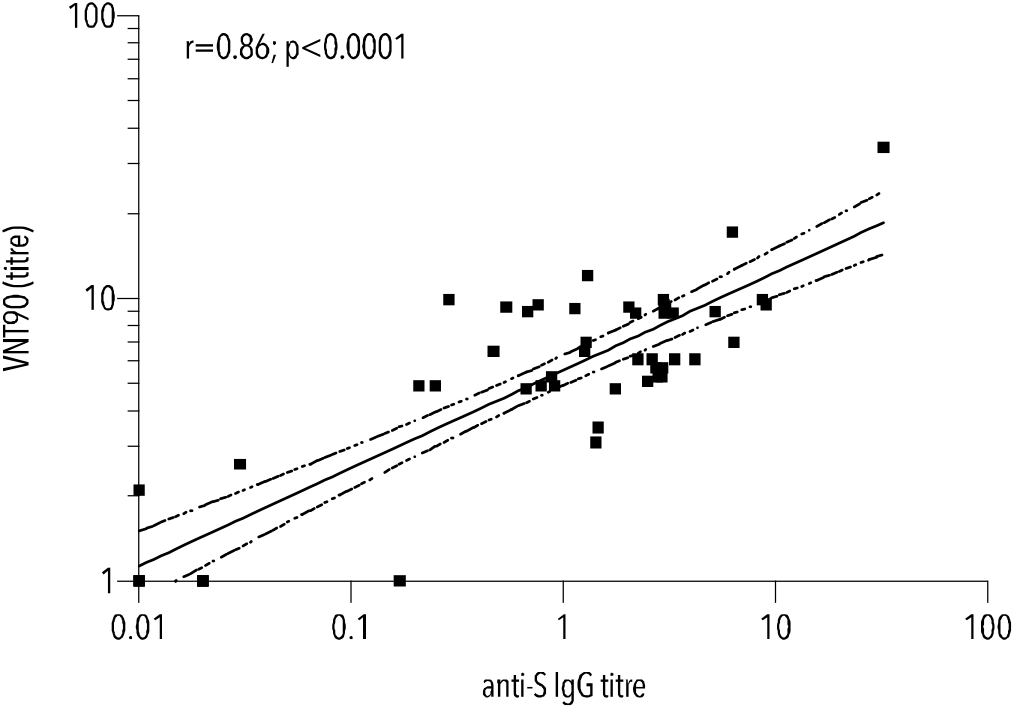
Correlation between anti-S IgG and VNT_90_ titers of antibodies (r … Pearson correlation coefficient; VNT_90_ … virus-neutralizing antibody titer inhibiting 90% of infectious dose)

Post-vaccination responses established by the change of selected lymphocytes populations in the murine spleens, i.e., plasmablasts, short- and long-lived plasma cells, including memory B cells, as well as Th1, Th2, Th17 and follicular Th lymphocytes, were observed in immunized mice compared with unimmunized ones regardless of the route of administration and number of doses received (Figure 6). While a significant increase was observed in follicular Th lymphocytes for both routes of administration compared with unimmunized mice, a borderline increase in memory B cells was seen only in mice with ID vaccination. No post-vaccination changes in the subsets of regulatory T cells or cytotoxic T lymphocytes as well as active B cells were found (data not shown).

**Figure 6:**
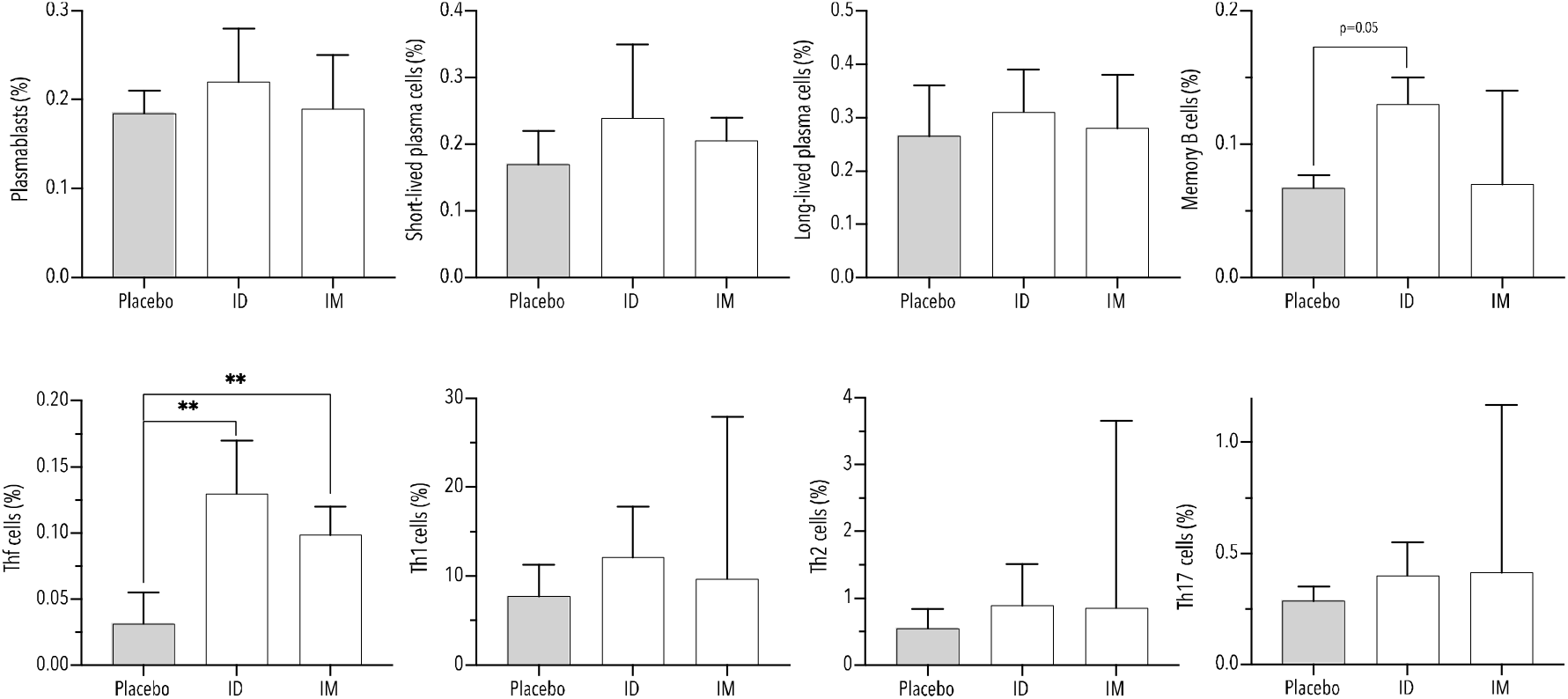
Median and range of merged subpopulation of B and T cells for mice immunized with ID or IM vaccine and placebo, irrespective of the size and number of doses. (ID … intradermal administration; IM … intramuscular administration; Thf … follicular helper T cells; * … p<0.05; ** … p<0.01)

The follicular Th lymphocytes demonstrated an upward trend depending on the time or number of received doses in immunized mice regardless of the dose size or route of administration (Figure 7B). The slope of the regression line differing significantly from null (p<0.05) revealed that mice immunized intradermally with a 0.5 μg dose exhibited significantly increasing proportions not only of follicular Th lymphocytes but also Th1 and Th2 lymphocytes with an increasing number of doses (Figure 7A). In addition, the same effect was observed in these mice for subsets of plasmablasts and memory B cells.

**Figure 7:**
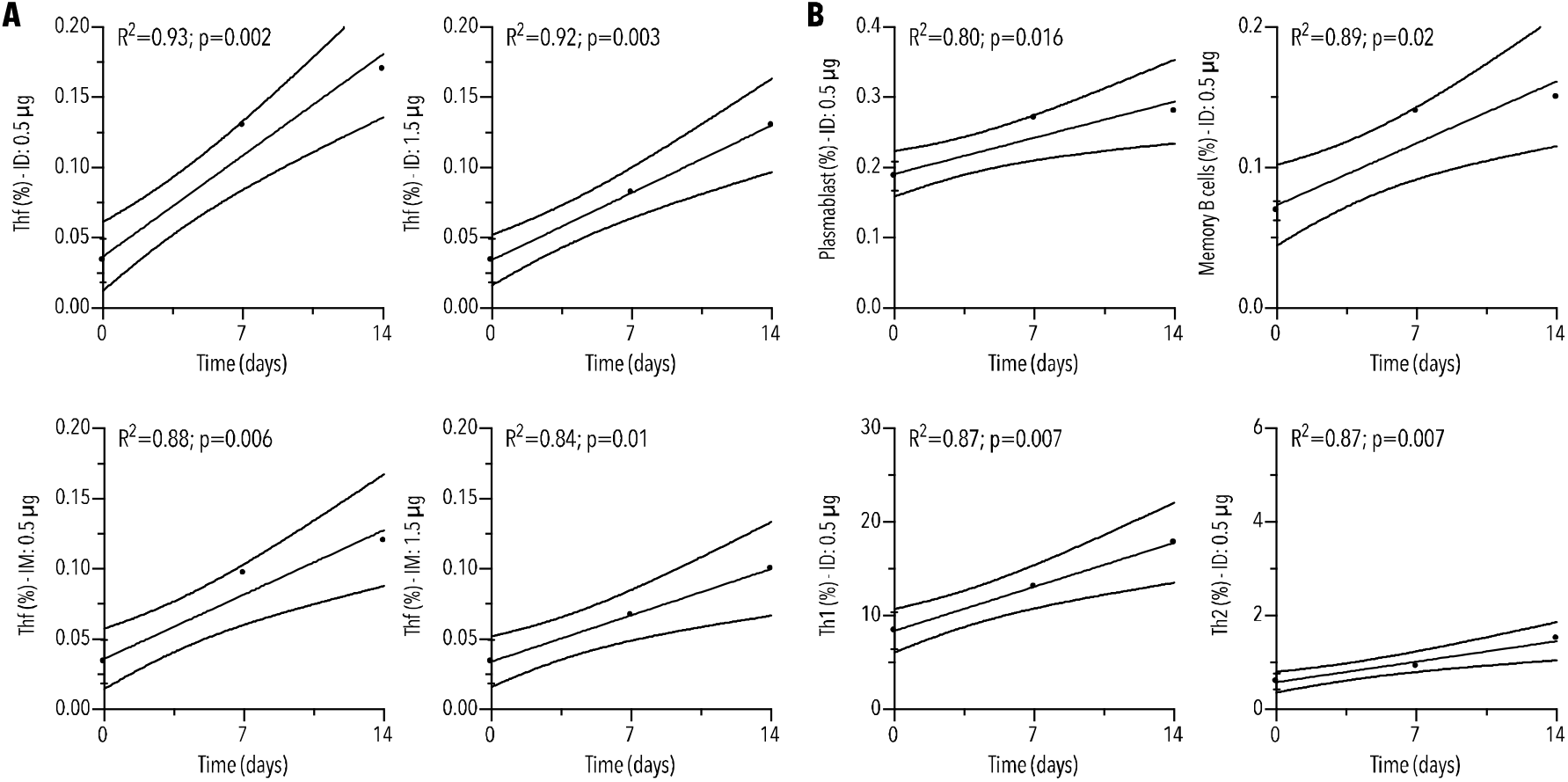
Linear regression of the B and T cells subsets exhibiting a non-null slope of the dose-number-dependent increase; A – follicular Th cells after both routes of administration and for both doses of 0.5 and 1.5 μg; B – plasmablasts, memory B cells, Th1 and Th2 cells after intradermal administration of the 0.5 μg dose (ID … intradermal administration; IM … intramuscular administration; Thf … follicular helper T cells; R^2^ … coefficient of determination)

## Discussion

The above laboratory procedure generated a semi-split inactivated vaccine, i.e., a vaccine with the S protein separated from the viral particle exhibiting an early, both humoral and cellular, immune response.

The design of this prototype vaccine was based on the usual procedures [Tang 2003, Spruth 2005, Gao 2020]. We selected a strain showing higher infectivity with lower virulence, i.e., one that showed signs of attenuation. Overall, we have tested several candidate strains for both immunogenicity through convalescent plasma and propagation capacity by viral titration.

Viral growth kinetics analysis was in line with the published data [Gao 2020, Manenti 2020]. Furthermore, we devised a technique to increase the virus concentration in suspension about 100 to 1,000-fold (note: this finding requires validation).

To inactivate the virus properly, beta-propiolactone was chosen because it can alkylate the viral genome and keeps the virus capable of inducing a protective immune response [Delrue 2012]. The process of viral suspension double inactivation was designed in accordance with the procedures of other vaccine manufacturers [Pu 2020, Xia 2020].

Use of the purification technique proposed by us resulted in an anticipated separation of S proteins from viral particles. We assumed that our semi-split vaccine could better offer all of the four important antigens to the immune system, i.e. spike, envelop, membrane and nucleocapsid proteins. The semi-split inactivated vaccine induced a humoral immune response as demonstrated by both anti-S IgG and virus-neutralizing antibodies.

Unfortunately, our immunogenicity results cannot be compared with the currently available data since we focused on monitoring the very early response, i.e., 7 days after one or two administered doses.

The success of this semi-split vaccine was a rapid onset of the immune response. It was slightly higher after intradermal than intramuscular administration independently of the number of doses and post-vaccination period. The second dose contributed to an increase in both anti-S IgG and virus-neutralizing antibody levels, but only neutralizing antibodies induced by two IM doses exhibited a significant rise compared with one IM dose regardless of size dose.

The post-vaccination cellular immunity suggested increased proportions of B and T lymphocytes subsets in the spleen such as plasmablasts, long- and short-lived plasma cells, memory B cells, Th1 and Th2 including Th17 lymphocytes whatever the route of administration, number and size of doses compared with the lymphocyte counts of placebo-treated mice. Obviously, a significant elevation was achieved in follicular Th lymphocytes for both intradermal and intramuscular vaccine administration.

Surprisingly, a marked upward trend in plasmablasts, memory B cells, Th1, Th2 and follicular Th lymphocytes was demonstrated in mice immunized intradermally with one and two 0.5 μg doses. It is not clear at the moment whether or not this effect is influenced by dose size or exclusively by the intradermal route of administration.

As we have evaluated only the early immune response to date, it cannot be concluded that a higher dose or intramuscular administration would be inferior in the final response. Still, early immunogenicity was better as demonstrated in mice with intradermal immunization.

We are well aware of the need to further investigate the other specific antibodies, i.e., not only against the S protein but, also, against the E, M and N proteins as potential candidates not tested in the current phase of development yet. The body of knowledge obtained from an immuneinformatics approach suggests that the design of a multi-epitope vaccine including all four antigens could hold promise to achieve efficient and safe vaccination against SARS-CoV-2 [Abdelmageed 2020].

## Conclusion

As it is, the semi-split inactivated vaccine seems to be capable of inducing both humoral and cellular immunity, especially after its intradermal administration. The early immune response, reasonable safety profile observed in the animal model as well as potential multi-antigenic extended post-vaccination protection are likely to be acceptable for human immunization.

