## Supplement for "Early immune response in mice immunized with a semi-split inactivated vaccine against SARS-CoV-2 containing S protein-free particles and subunit S protein": Supplement.docx

Deep immunophenotyping of splenocytes using spectral flow cytometry

As shown in Table 1, a 24-color spectral cytometry panel was designed to characterize T and B cell differentiation and activation status.

Briefly, cryopreserved splenocytes were thawed and washed with RPMI 1640 supplemented with 10% fetal bovine serum (FBS), 0.2% bicarbonate, 1 mg/l folic acid, 0.4 mM HEPES, 50 µM 2-mercapto-ethanol, 2 mM glutamine and the antibiotics penicillin and streptomycin (Gibco, USA). The cells were subsequently resuspended in supplemented RPMI 1640 and rested overnight to normalize marker expression after freezing. After resting, cells were counted and up to 4 million cells per sample were used for further staining. In the first step, the cells were washed with phosphate buffer saline (PBS) to remove medium residua and subsequently stained with Live Dead Blue dye (Thermo Fisher, Waltham, MA) at room temperature for 20 min. After that, the cells were washed using FACS buffer (PBS with 2% BSA, 0.1% NaN_3_) and Fc receptors were blocked with Trustain FcX (anti-mouse CD16/32) antibody (Biolegend, San Diego, CA). Next, the cells were stained with a combination of surface antibodies (as listed in Table 1S) at room temperature for 30 min. Following surface staining, the cells were washed with FACS buffer, fixed and permeabilized using FoxP3/Transcription Factor Staining Buffer Set (Thermo Fisher). Finally, the cells were stained with an anti-FoxP3 antibody (Table 1S). A minimum of 1.5 million cells were acquired using the Aurora spectral cytometer (Cytek Biosciences, Fremont, CA) equipped with 5 lasers (355, 405, 488, 561, 640 nm) and 64 detectors.

Data Analysis

The acquired data were unmixed and manual analysis of data was performed using FlowJo 10.7.1 software (BD Biosciences, Ashland, OR). The gating strategy used for data analysis is shown on Figure 1S. For unsupervised clustering analysis all fcs files were pre-processed by removing debris, doublets and dead cells. Then, these files were biexponentially transformed and used to train EmbedSOM [Kratochvíl 2020] with 32x32 SOM grid.

| Table 1S. Reagents used for polychromatic flow cytometry | | | | |
| --- | --- | --- | --- | --- |
| Name | Fluorochrome | Clone | Manufacturer | Staining |
| CD3 | Spark Blue 550 | 17A2 | Biolegend | surface |
| CD335 (Nkp46) | PE/Dazzle 594 | 29A1.4 | Biolegend | surface |
| CD4 | PerCP | RM4-5 | BD Biosciences | surface |
| CD25 | PeCy7 | PC61 | Biolegend | surface |
| CD19 | BV570 | 6D5 | Biolegend | surface |
| CD8 | BV650 | 53-6.7 | Biolegend | surface |
| CD184 (CXCR4) | BV786 | 2B11 | BD Biosciences | surface |
| FoxP3 | PE | MF-14 | Biolegend | intracellular |
| IgD | Pacific Blue | 11-26c.2a | Biolegend | surface |
| CD196 (CCR6) | BV421 | 29-2L17 | Biolegend | surface |
| CD62L | BUV805 | MEL-14 | BD Biosciences | surface |
| CD185 | BUV737 | 2G8 | BD Biosciences | surface |
| CD93 | BUV615 | AA4.1 | BD Biosciences | surface |
| IgM | PerCPCy5.5 | R6-60.2 | BD Biosciences | surface |
| CD27 | BUV496 | LG3A10 | BD Biosciences | surface |
| CD44 | BUV395 | IM7 | BD Biosciences | surface |
| CD45R (B220) | BB700 | B220 | BD Biosciences | surface |
| CD138 | BB515 | 281-2 | BD Biosciences | surface |
| CD38 | APC/Cy7 | 90 | Biolegend | surface |
| CD194 (CCR4) | APC | 2G12 | Biolegend | surface |
| CD183 (CXCR3) | BV480 | CXCR3-173 | BD Biosciences | surface |
| I-A/I-E | Alexa Fluor 700 | M5/114.15.2 | Biolegend | surface |
| CD279 (PD-1) | BV711 | 29F.1A12 | Biolegend | surface |
| Viability Amine Reactive Dye | Live Dead Blue | - | Thermo Fisher | Viability |

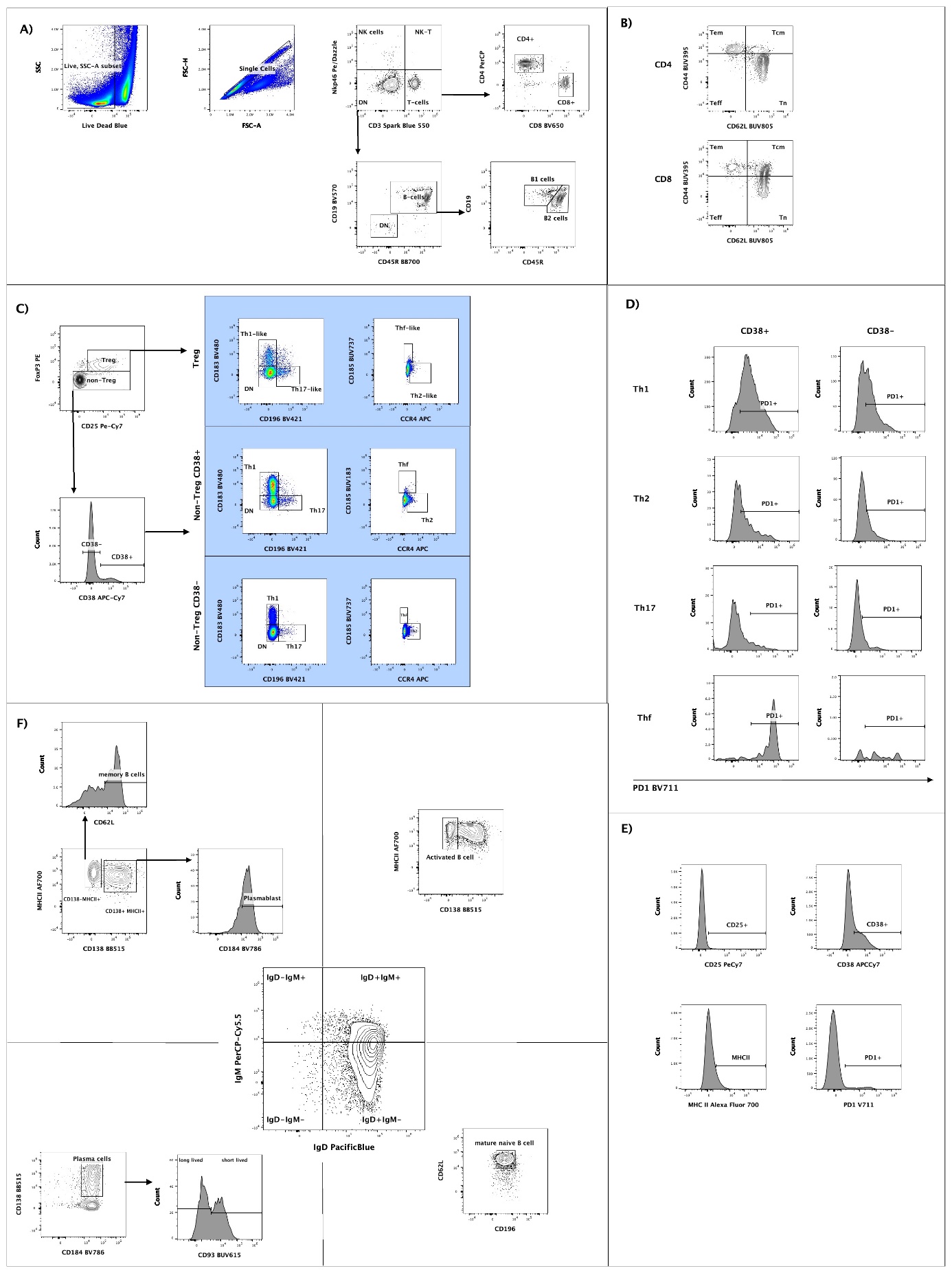

Figure 1S: Gating strategy for flow cytometry data analysis

**A)** Major lymphocyte lineages were identified by key markers: CD3 for T cells, CD335 (Nkp46) for NK cells and finally B cells were defined based on the expression of CD19 and CD45R (B220). T cells were subsequently subdivided into helper T cells (Th) and cytotoxic T cells according to CD4 and CD8 expression. B cells were subdivided into CD19+CD45R^low^ B1 cells that are considered to be part of ^the^ innate immune system, and classical CD19+CD45R+ B2 cells [Montecino-Rodriguez, Leathers, and Dorshkind 2006].
 **B)** In both CD4+ and CD8+ T cell subsets, several populations can be defined based on the expression of CD44 and CD62L. The CD44+CD62L- population consists of effector memory cells (Tem), CD44+CD62L+ correspond to central memory cells (Tcm), CD44-CD62L+ represent naïve T cells (Tn) and, finally, CD44-CD62L- define effector cells (Ahlers and Belyakov 2010). **C)** The CD4+ Th population consists of several subtypes with different functional properties. Regulatory T cells (Treg) are responsible for controlling inflammation and peripheral tolerance. They are defined based on the expression of CD25, the high affinity receptor for IL2, and FoxP3. Tregs can be further divided based the on expression of chemokine receptors into Th-like subsets.

Activation status of T cells was assessed based on the expression of the activation marker CD38; this allows for the identification of Th subsets that had been activated by antigen stimuli. There are several non-Treg CD4+ T cell subsets. Th1 cell are responsible for antiviral immunity and are the main source of IFNγ; these are defined based on CD183 expression. Th17 cells are the main drivers of anti-bacterial and anti-fungal immunity and are defined based on the expression of CD196. Th2 and follicular helper T cells responsible for regulating antibody responses can be identified based as CD183-CD196-CD185-CCR4+ and CD183-CD196-CD185+CCR4-, respectively [Sandoval-Montes and Santos-Argumedo 2005; Stubbington et al. 2015] .

**D)** Expression of coinhibitory receptors such as PD1 can be induced based on the strength of antigen stimulation and used as a marker of T cell exhaustion, except in Thf cells which express PD1 constitutively.

**E)** Cytotoxic CD8+ T cells are responsible for elimination of virus-infected cells. Their activation was assessed using expression of activation markers such CD25, MHCII, CD38; furthermore PD1 was monitored as a marker of exhaustion.

**F)** The B2 cell subset was further analyzed to identify key subsets and monitor the respective changes in these after vaccination. Phenotyping of the B2 cell compartment was primary based on the expression of IgM and IgD. Mature naïve B cells that have not reacted did not react with antigen were defined as IgM-IgD+CD62L+CD196+. Activated B cells were characterized as IgM+IgD+MHCII+CD138-. In the IgM+IgD+ compartment, 2 subsets of B cells were defined, IgM+IgD+MHCII+CD138-CD62L+ memory B cells and IgM+IgD+MHCII+CD138+CD184+ plasmablasts. Finally, antibody-secreting plasma cells were defined as IgM-IgD-CD138+CD184+ cells. These were further divided based on the expression of CD93 into CD93- long-lived and CD93- short-lived CD93+ plasma cells [Sarvaria, Madrigal, and Saudemont 2017; Shen and Fillatreau 2015; Suan et al. 2017].

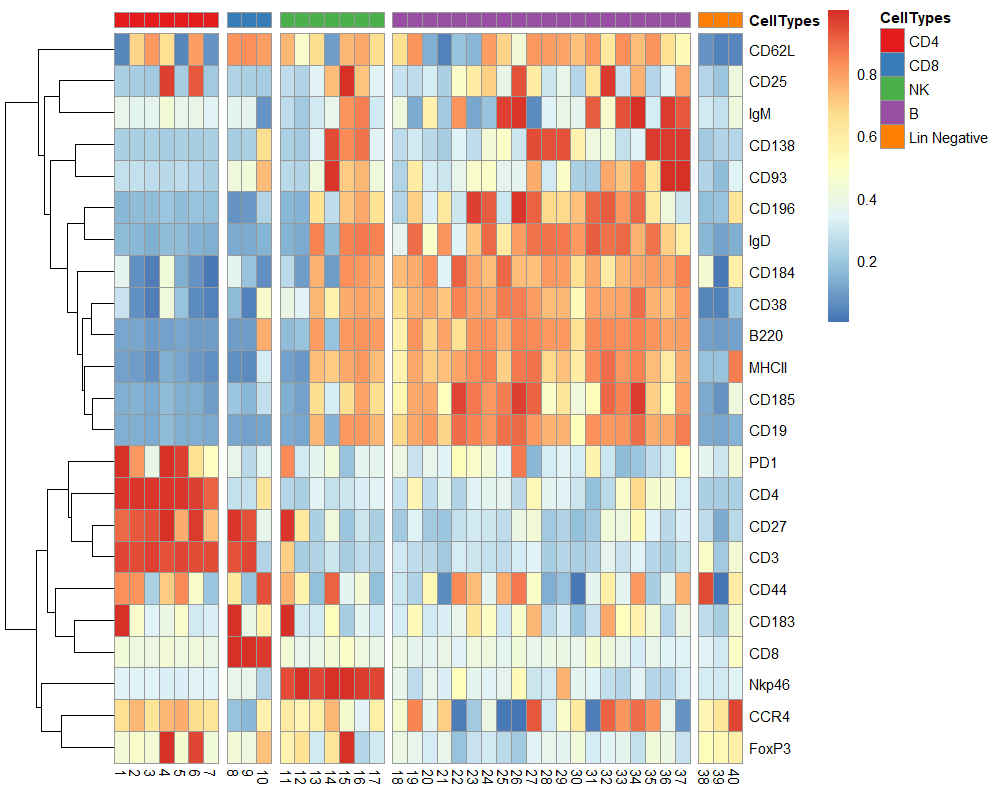

**C)**

**B)**

**A)**

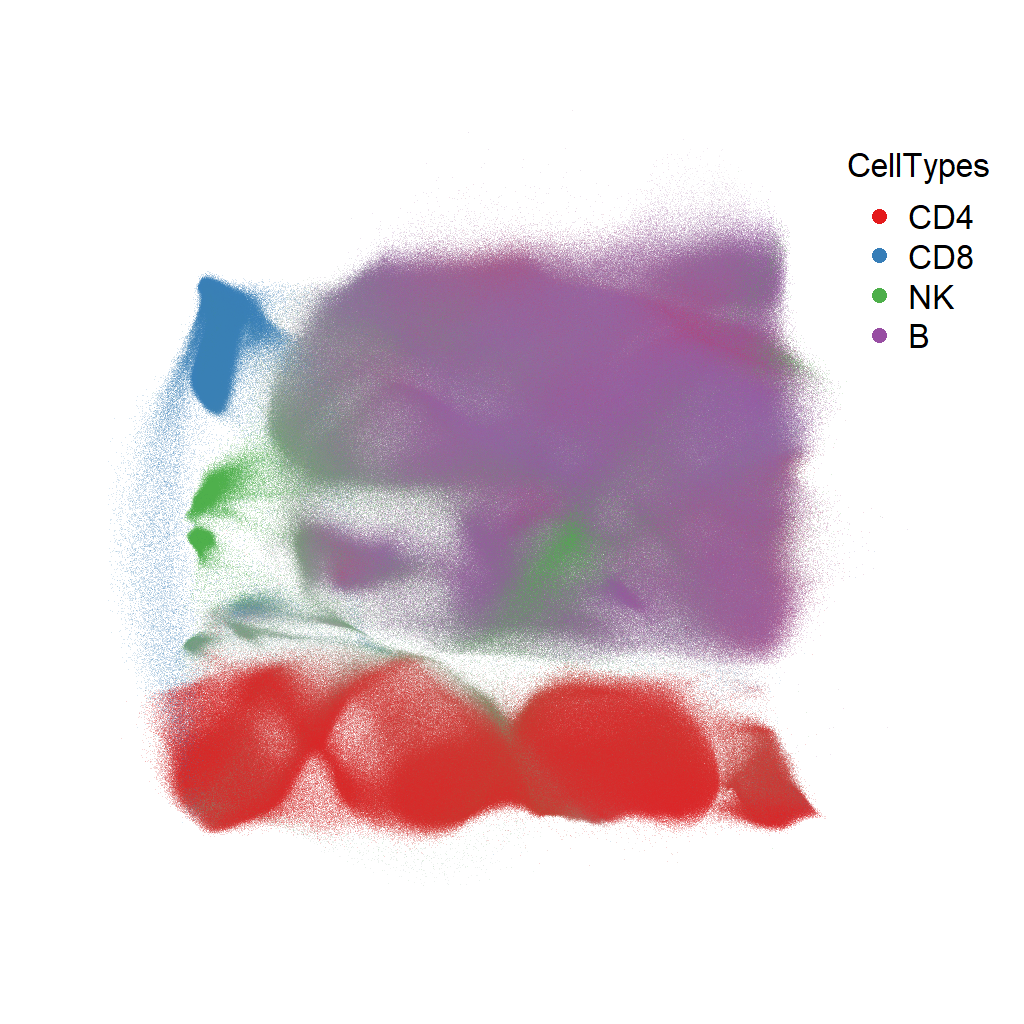

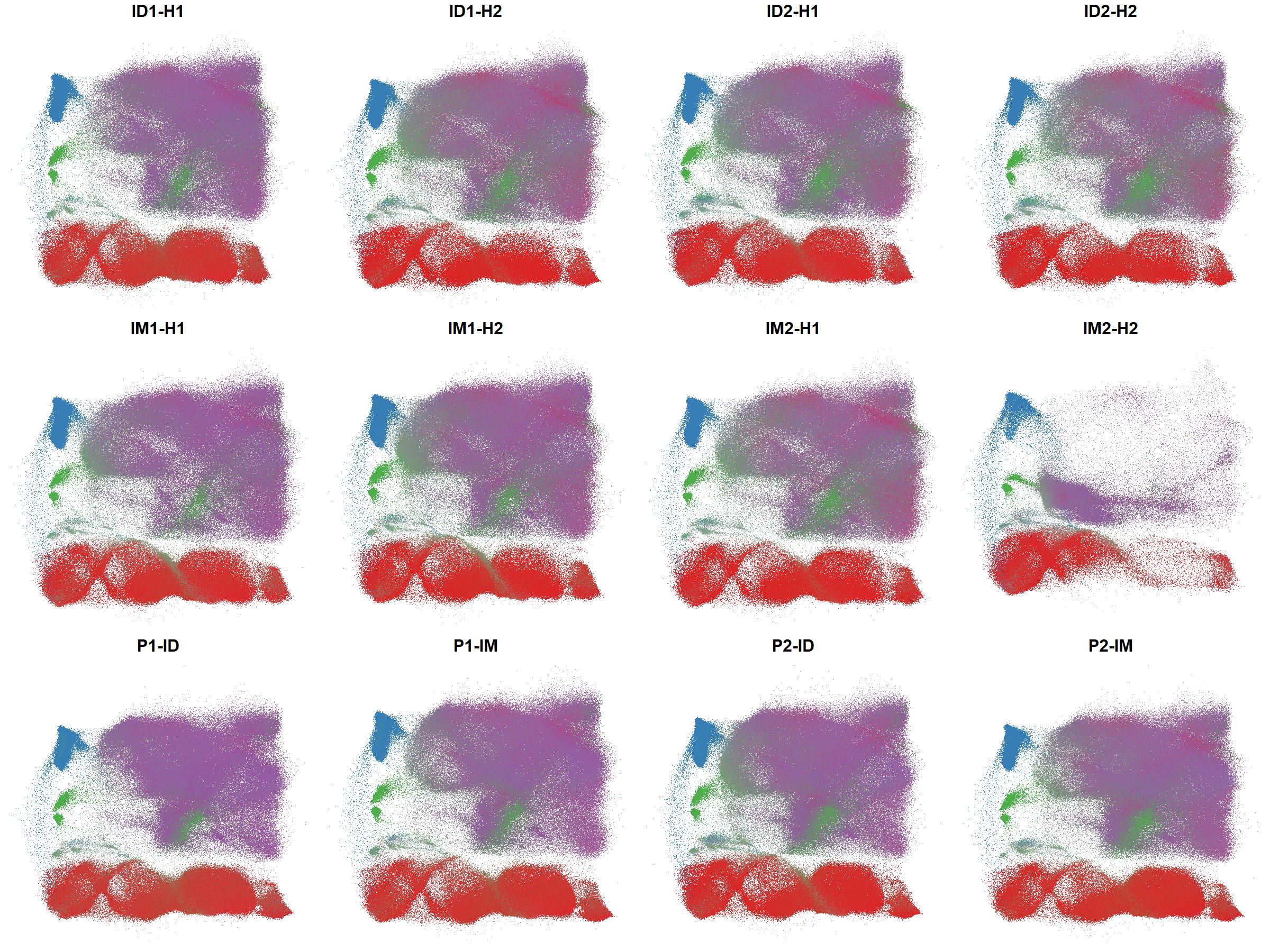

Figure 2S: Unsupervised clustering analysis of major cell types and their characterization using EmbedSOM algorithm

**A)** EmbedSOM representation of cellular lineages of T, NK, B cells from all fcs files and

**B)** individual files (ID- Intra dermal, IM- Intra Muscular, P- Placebo, H1- Harvest-1, H2-Harverst-2).

**C)** Heatmap representation of 40 meta-clusters identified by EmbedSOM. The expression of individual cluster was correlated with manual gating based on markers of expression. For CD4 T cells total 7 cluster were identified out of which cluster 4 and 7 represents T-regs, cluster 1 represents Th1, cluster 5 Th2 and cluster 2 represents Thf cells. Where as in B cells compartment clusters 19,21,23,24,26,27,31 to 37 are B1 cells whereas the rest are B2 cells. .

Supplementary references

Ahlers, Jeffrey D., and Igor M. Belyakov. 2010. “Memories That Last Forever: Strategies for Optimizing Vaccine T-Cell Memory.” *Blood*.

Kratochvíl, Miroslav, Abhishek Koladiya, and Jiří Vondrášek. 2020. “Generalized EmbedSOM on Quadtree-Structured Self-Organizing Maps.” *F1000Research*.

Montecino-Rodriguez, Encarnacion, Hyosuk Leathers, and Kenneth Dorshkind. 2006. “Identification of a B-1 B Cell-Specified Progenitor.” *Nature Immunology*.

Sandoval-Montes, Claudia, and Leopoldo Santos-Argumedo. 2005. “CD38 Is Expressed Selectively during the Activation of a Subset of Mature T Cells with Reduced Proliferation but Improved Potential to Produce Cytokines.” *Journal of Leukocyte Biology*.

Sarvaria, Anushruti, J. Alejandro Madrigal, and Aurore Saudemont. 2017. “B Cell Regulation in Cancer and Anti-Tumor Immunity.” *Cellular and Molecular Immunology*.

Shen, Ping, and Simon Fillatreau. 2015. “Antibody-Independent Functions of B Cells: A Focus on Cytokines.” *Nature Reviews Immunology*.

Stubbington, Michael J.T. et al. 2015. “An Atlas of Mouse CD4+ T Cell Transcriptomes.” *Biology Direct*.

Suan, Dan et al. 2017. “CCR6 Defines Memory B Cell Precursors in Mouse and Human Germinal Centers, Revealing Light-Zone Location and Predominant Low Antigen Affinity.” *Immunity*.
